# Loss of Cathepsin K impairs collagen biogenesis and enhances actin polymerization in trabecular meshwork

**DOI:** 10.1101/2025.02.10.637394

**Authors:** Avinash Soundararajan, Krishna Jaysankar, Emma Doud, Rodahina Philihina Pasteurin, Michelle Surma, Padmanabhan P Pattabiraman

**Affiliations:** Ophthalmology, Indiana University School of Medicine, Indianapolis, Indiana, United States; Medical Neuroscience Graduate Program, Stark Neuroscience Research Institute, Indiana University School of Medicine, Indianapolis, Indiana, USA

**Keywords:** Cathepsin K, extracellular matrix, actin cytoskeleton, intraocular pressure, trabecular meshwork

## Abstract

Trabecular meshwork (TM) dysfunction and extracellular matrix (ECM) dysregulation contribute to increased intraocular pressure (IOP) in primary open-angle glaucoma (POAG). Earlier, we provide a proof-of-concept study identifying the regulation and the role of Cathepsin K (CTSK), a potent collagenase, in ECM homeostasis, actin bundling, and IOP regulation. Better understanding of the loss of CTSK function in TM remains unclear. Using siRNA-mediated knockdown of CTSK (siCTSK) in human TM cells, this study investigated the role of CTSK in actin and ECM homeostasis using an unbiased proteomics approach. Loss of CTSK significantly disrupted collagen biogenesis and ECM homeostasis. CTSK depletion also increased intracellular calcium levels, with proteomics data suggesting possible involvement of calcium-regulatory proteins. Additionally, PRKD1 activation enhanced actin polymerization through the LIMK1/SSH1/cofilin pathway, promoting focal adhesion maturation. Despite increased apoptotic markers (CASP3, CASP7, TRADD, PPM1F), caspase 3/7 activation was not induced, suggesting apoptosis-independent cellular remodeling. Notably, RhoQ and myosin motor proteins were significantly downregulated, indicating altered mechanotransduction in TM cells. These findings highlight the role of CTSK in maintaining ECM homeostasis, calcium signaling, and cytoskeletal regulation in TM. Its depletion induces actin polymerization, which may influence aqueous humor outflow. Targeting CTSK-related pathways may provide novel therapeutic strategies for regulating IOP and preventing glaucoma progression.

## 1. Introduction

Glaucoma is a progressive optic neuropathy and the leading cause of irreversible blindness worldwide (1). In 2020, approximately 76 million people were estimated to be affected by glaucoma, with this number projected to rise to 111.8 million by 2040 (1). Of these, primary open-angle glaucoma (POAG) accounts for majority of cases (2). The primary risk factor for developing POAG is elevated intraocular pressure (IOP), the only modifiable factor for slowing the disease’s progression (3). Aqueous humor, a fluid that nourishes the eye, is continuously produced by the ciliary body and drained primarily through the trabecular meshwork (TM). This balance between production and drainage regulates IOP within the eye (4). The TM is a specialized tissue composed of extracellular matrix (ECM) components, including collagens, elastin, and glycosaminoglycans, which provide structural integrity and elasticity for aqueous humor outflow (5). The ECM, along with the actin cytoskeleton in TM cells, plays a critical role in regulating aqueous humor drainage and maintaining IOP (6). Proper drainage is dependent on a finely tuned balance of ECM homeostasis, which involves the synthesis and degradation of ECM components (7). This balance is orchestrated by a variety of enzymes, including matrix metalloproteinases (MMPs) and cathepsins, which mediate ECM remodeling (8, 9). Dysregulation of ECM homeostasis, such as excessive ECM deposition or inefficient degradation, can lead to impaired aqueous humor outflow, contributing to increased IOP and the pathogenesis of glaucoma (10).

Cathepsin K (CTSK) is a cysteine protease that is primarily known for its critical role in ECM degradation, particularly type 1 collagen (11). Our previous study demonstrated that CTSK is negatively regulated by pathological stressors that disrupt ECM remodeling in the TM and contribute to IOP dysregulation (9). We further reported that pharmacological inhibition of CTSK activity using balicatib resulted in increased collagen deposition and actin bundling in TM cells, leading to elevated IOP. Conversely, induction of CTSK expression reduced ECM accumulation and promoted actin depolymerization through the dephosphorylation of cofilin (9).

The mechanistic role of CTSK in modulating ECM components within the TM remains poorly understood, representing a critical gap in our knowledge. Understanding these mechanisms is essential, as they have significant implications for IOP regulation and glaucoma pathogenesis. In this study, we sought to elucidate the molecular pathways regulated by CTSK in TM cells by performing a global proteomic analysis following siRNA-mediated CTSK knockdown. This approach enabled us to identify differentially expressed proteins and pathways associated with ECM remodeling, cytoskeletal dynamics, and cellular signaling in the absence of CTSK in TM.

Our findings provide a foundation for advancing therapeutic strategies targeting CTSK to better manage ocular hypertension and primary open-angle glaucoma.

## 2. Materials and methods

### 2.1 Materials

The reagents and antibodies used in this study are presented in **Table 1**.

**Table 1:**
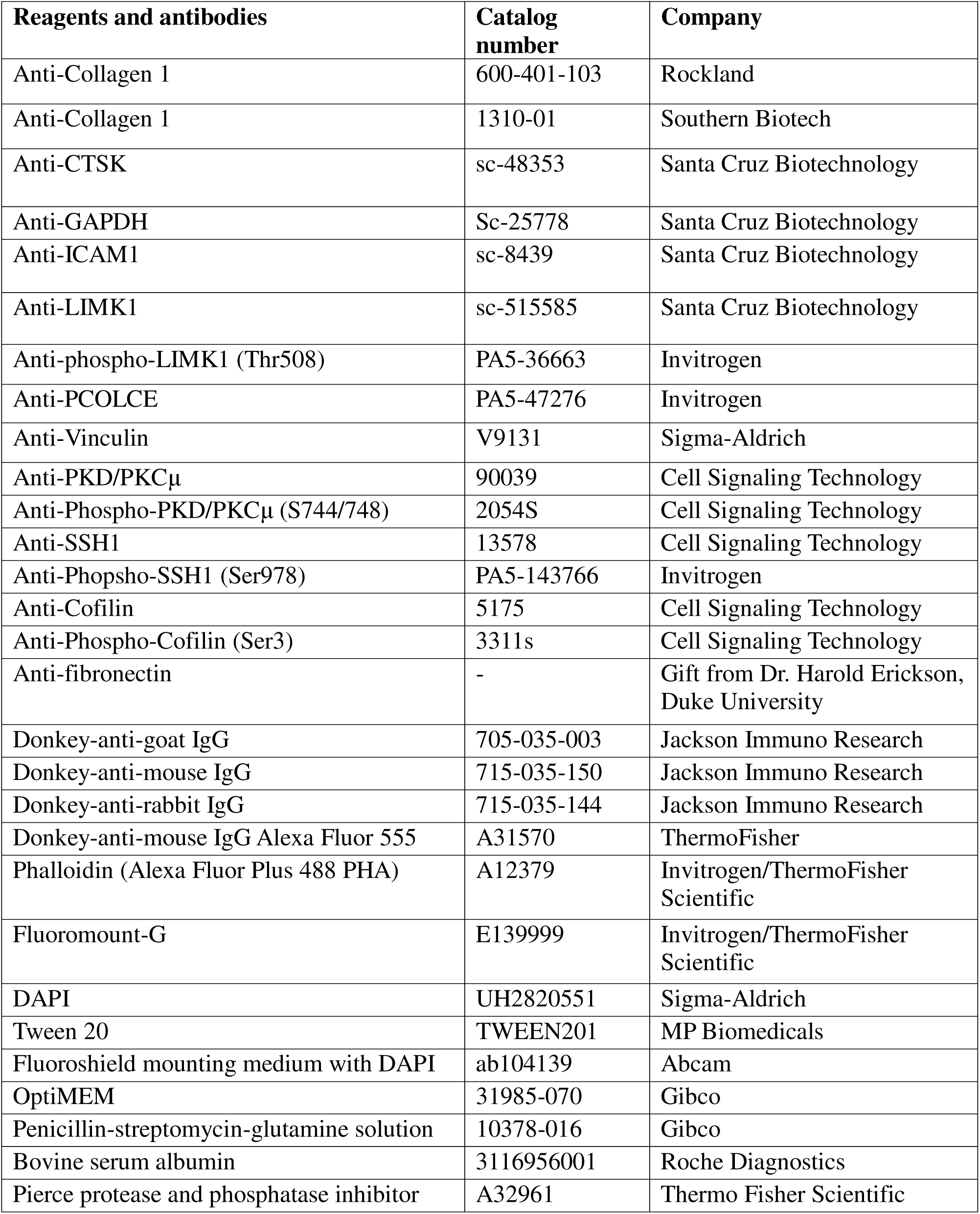
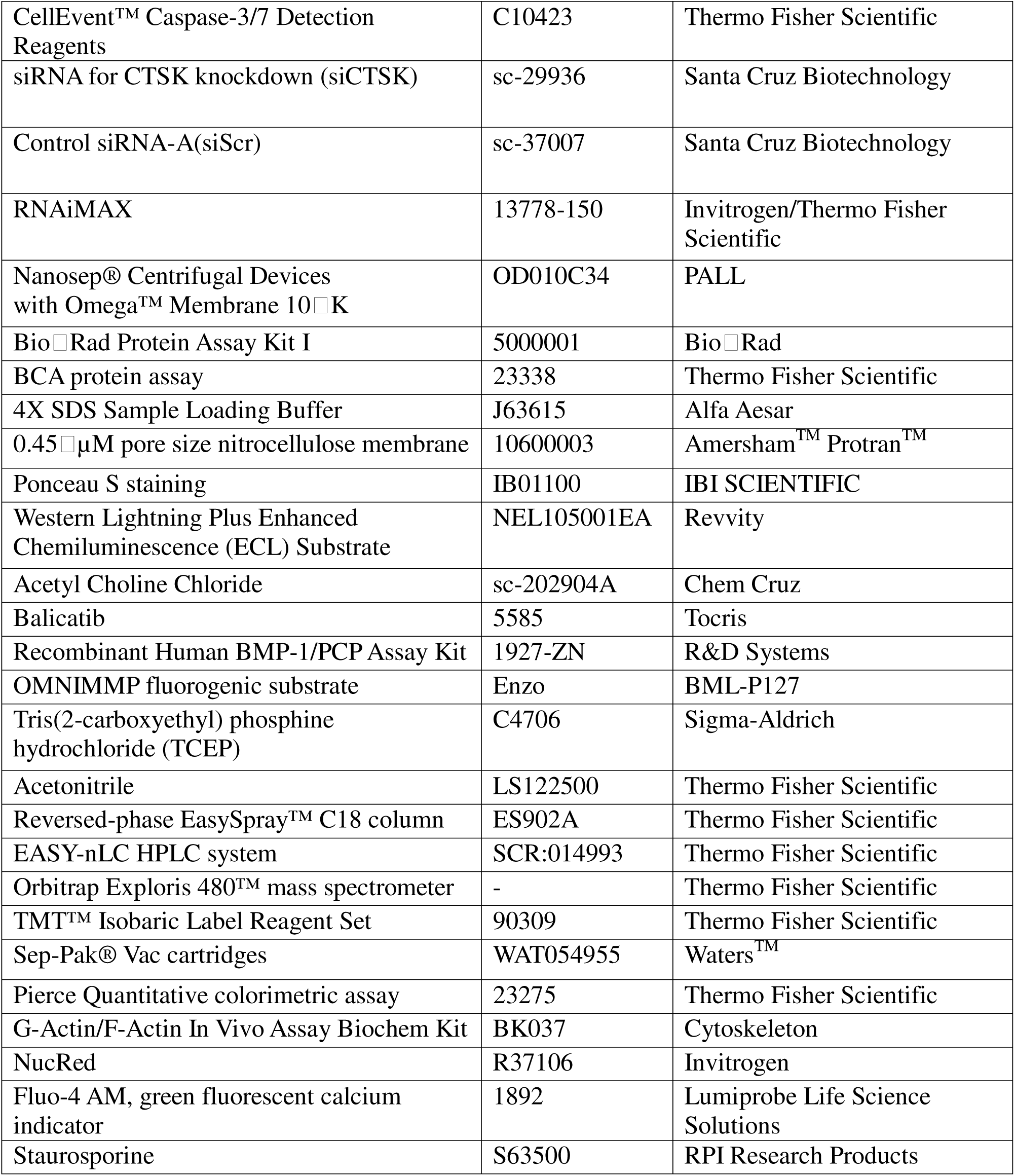
Materials and reagents used.

### 2.2 Primary trabecular meshwork cell culture

Primary human TM (HTM) cells were cultured from TM tissue isolated from the leftover donor corneal rings after they had been used for corneal transplantation at the Indiana University Clinical Service, Indianapolis, and characterized as described previously (12). HIPPA compliance guidelines were adhered for the use of human tissues. The usage of donor tissues was exempt from the DHHS regulation and the IRB protocol (1911117637), approved by the Indiana University School of Medicine IRB review board. Donor age, race, and sex details are provided in **supplementary table 1**. TM tissue extracted from the corneal ring was chopped into fine pieces and placed in a 2% gelatin-coated six well tissue culture plate sandwiched by a coverslip. The tissues were grown in OptiMEM, containing 20% fetal bovine serum (FBS) and penicillin/streptomycin/glutamine solution. The expanded population of HTM cells was sub cultured after 1–2 weeks in Dulbecco’s modified Eagle’s medium, containing 10% FBS and characterized by the detection of dexamethasone-induced myocilin (10, 13). For our experiments, we used confluent cultures between the fourth and sixth passages. All experiments in this manuscript were performed using biological replicates.

### 2.3 siRNA mediated knockdown of CTSK

To investigate the loss of CTSK function in TM cells in vitro, we performed siRNA-mediated knockdown of CTSK. Human trabecular meshwork (HTM) cells were transfected with either scramble control siRNA (siScr) or CTSK-specific siRNA (siCTSK) using RNAiMAX transfection reagent. Transfections were carried out in Opti-MEM. After 12 hours of incubation, the medium was replaced with serum-free DMEM. Seventy-two hours post-transfection, cells were harvested in 8M urea buffer for proteomic analysis or in RIPA buffer for immunoblotting. Additionally, the culture media was concentrated using Nanosep® centrifugal devices with a 10K Omega™ membrane.

### 2.4 Protein expression analysis by immunoblotting

The cell lysates containing total protein were prepared using 1X RIPA buffer composed of 50 mM Tris HCl (pH 7.2), 150 mM NaCl, 1% NP-40, 0.1% sodium dodecyl sulfate (SDS), 1 mM EDTA, and 1 mM PMSF with protease and phosphatase inhibitors and then sonicated. The protein concentration was determined using either the Bradford Assay Reagent/Bio-Rad Protein Assay Kit I or BCA protein assay. A total of 20–40 μg of the protein sample was mixed with 4X Laemmli buffer and separated on 8%–15% SDS polyacrylamide gel. Following the gel run, the proteins were transferred to 0.45 μm pore size nitrocellulose membrane. Ponceau S staining of the membrane was performed to document the protein loading after transfer. Membranes were blocked in 5% nonfat dry milk in Tris buffered saline with 0.1% Tween for 2 h followed by respective primary antibodies overnight at 4°C (∼16 h), and then by horseradish peroxidase-conjugated secondary antibodies. The blots were washed with 1X Tris buffered saline with Tween-20, and the immunoreactivity was detected using Western Lightning Plus Enhanced Chemiluminescence (ECL) Substrate and imaged using ChemiDoc MP imaging system (Bio Rad). Blots were stripped using mild stripping buffer if required to re-probe for the loading control and multiple proteins within the same molecular weight range. The data were normalized to GAPDH or β-actin. Semi-quantitative analyses and fold changes were calculated from the band intensities measured using Image J software.

### 2.5 Protein distribution analysis by immunofluorescence

HTM cells were cultured on glass coverslips coated with 2% gelatin until they reached 90% confluency. Following the necessary treatments, the cells were rinsed twice with 1X PBS, then fixed with 4% paraformaldehyde for 15 minutes. The cells were permeabilized using 0.2% Triton X-100 in PBS for 10 minutes and subsequently blocked with 5% bovine serum albumin in 1X PBS for 1 hour. The cells were then exposed to the primary antibody and incubated overnight at 4°C. After two washes with 1X PBS, the cells were incubated with Alexa Fluor-conjugated secondary antibodies for 2 hours at room temperature. Finally, the coverslips were rinsed and mounted on glass slides using Fluoroshield with DAPI Mounting Medium. Confocal microscopy was performed with a Zeiss LSM 700, and z-stack images were captured and processed using Zeiss ZEN software.

### 2.6 Global proteomics analysis Sample preparation

For proteomics analysis, cell pellets or ECM-enriched samples were subjected to lysis in 8 M Urea with 100 mM Tris-HCl (pH 8.5) via sonication using the Bioruptor® Plus sonicator. The samples were sonicated in 1.5 ml microtubes under cycles of 30 seconds on and 30 seconds off for a total of 15 minutes in a water bath kept at 4°C. After lysis, the lysates were centrifuged at 14,000g for 20 minutes, and the protein concentration was determined using the Bradford assay. Twenty micrograms of protein from each sample were treated with 5 mM tris(2-carboxyethyl) phosphine hydrochloride (TCEP) at room temperature for 30 minutes to reduce disulfide bonds. The exposed cysteine thiols were alkylated by incubation with 10 mM chloroacetamide at room temperature in the dark for 30 minutes. The samples were then diluted with 50 mM Tris-HCl (pH 8.5) to reduce the urea concentration to 2 M and digested overnight at 37°C using Trypsin/LysC in a protease-to-substrate ratio of 1:100.

#### Peptide purification and TMT labeling

Following digestion, peptides were acidified with 0.5% trifluoroacetic acid (TFA) and desalted using Sep-Pak® Vac cartridges. After a wash step with 0.1% TFA, the peptides were eluted in a 70% acetonitrile and 0.1% formic acid solution, then dried using a speed vacuum. The dried peptides were resuspended in 29 µL of 50 mM triethylammonium bicarbonate, and peptide concentrations were measured using the Pierce Quantitative colorimetric assay. For labeling, equal peptide amounts from each sample were reacted with 0.2 mg of Tandem Mass Tag (TMT™ Isobaric Label Reagent Set) at room temperature for 2 hours. The reactions were quenched with 0.3% hydroxylamine for 15 minutes at room temperature, and the labeled peptides were combined and dried using a speed vacuum.

#### Peptide fractionation and Nano-LC-MS/MS

The combined peptides were reconstituted in 0.1% TFA and fractionated using the Pierce™ High pH reversed-phase peptide fractionation kit. For Nano-LC-MS/MS analysis, the peptides were loaded onto a reversed-phase EasySpray™ C18 column and analyzed using a Nano-LC system connected to an Orbitrap Exploris 480™ mass spectrometer with a FAIMS pro interface. Approximately 1/8 of each global peptide fraction and 1/4 of each phosphopeptide fraction were loaded onto the column at 400 nl/min. The elution gradient started from 4% to 30% mobile phase B (0.1% FA, 80% acetonitrile) over 160 minutes, increased to 80% B over 10 minutes, and then decreased to 10% B over the final 10 minutes. The mass spectrometer was operated in positive ion mode using a data-dependent acquisition method with a 4-second cycle time and a FAIMS CV set at −50 V. Precursor ions were scanned across an m/z range of 375-1600 with a resolution of 60,000, and MS2 scans were conducted with a quadrupole isolation window of 0.7 m/z, utilizing 35% HCD CE at a resolution of 45,000.

#### Data processing and analysis

The RAW files generated from the mass spectrometry were processed using Proteome Discover™ 2.4 software. SEQUEST HT searches were conducted against a Homo sapiens UniProt FASTA database, including common contaminants, with a maximum of two missed cleavages permitted. The precursor mass tolerance was set at 10 ppm, and fragment mass tolerance at 0.02 Da. Static modifications incorporated carbamidomethylation on cysteine residues and TMT 6 plex labeling on N-termini and lysine residues, while dynamic modifications included oxidation of methionines and various modifications at protein N-termini. The false discovery rate (FDR) was controlled at 0.01 for stringent analysis and 0.05 for relaxed conditions. Normalization of peptides was based on total peptide amounts, with set thresholds for co-isolation and reporter ion signal-to-noise ratios. The resulting normalized values, fold changes, log2 ratios, and p-values were exported to Microsoft Excel for further analysis.

### 2.8 BMP1 assay

The PCP/BMP1 assay was performed following the manufacturer’s protocol. An assay buffer consisting of 25 mM HEPES with 0.1% Brij35 at pH 7.5 was used. Conditioned media from siCTSK/siScr or 10 μM balicatib was used for the assay. Recombinant BMP1, diluted in assay buffer, served as the positive control. The OMNIMMP fluorogenic substrate was added to each reaction. Fluorescence measurements were taken using a microplate reader at excitation and emission wavelengths of 320 nm and 405 nm, respectively.

### 2.9 Assessment of MMP activity using zymography

After treating HTM cells with siCTSK or siScr, the conditioned media was collected and concentrated. Equal volume of media was mixed with 2X non-reduced Laemmli sample buffer and separated by SDS-polyacrylamide gel electrophoresis containing 0.2% gelatin. The gel was renatured using 2.5% Triton X-100. Gels were incubated overnight (16–18 h) at 37°C in 50 mM Tris-HCl [pH 7.5], 200 mM NaCl, 5 mM CaCl2, 0.05% NaN3. The gel staining was carried out using 0.1% (w/v) Coomassie blue in staining/destaining buffer (Water: acetic acid: methanol-4.5:1:4.5) for 1.5 h. Destaining was performed using staining/destaining solution for approximately 2 h or until white bands are observed in clear background, which was imaged using ChemiDoc MP imaging system (Bio-Rad).

### 2.10 Flow cytometry

Apoptotic cell detection was performed using flow cytometry. Cells were treated with siSCR and siCTSK for 72 hours, with Staurosporine (STS) treatment serving as a positive control for apoptosis induction. Following treatment, both living and dead cells were collected and stained with NucRed and CellEvent™ Caspase 3/7 Green Detection Reagent. This reagent was diluted to a 5 μM final concentration in 1 mL of media. The diluted reagent was applied to the cells, which were then incubated for 2 hours at 37°C. Post-incubation, single-cell suspensions in PBS were run through the Attune NxT flow cytometer (Thermo Fisher) where 100 μL of cells was analyzed at a rate of 200μL/min. Cell debris was gated from a total cell population, and then the far-red positive (nucleus) and green positive (caspase) population was gated based on unstained cells. Fold changes of the double positive populations in siCTSK were compared to the control siSCR double positive population.

### 2.11 Calcium assay

Intracellular calcium levels were measured using Fluo-4 AM, a cell-permeable Ca^2+^ indicator, following the manufacturer’s instructions and previous studies (14, 15). After treatment, cells were incubated with 3 μM Fluo-4 AM at 37°C for 30 minutes and then washed with 1X PBS, followed by imaging using end point reading in Biotek Synergy H1 microplate reader and confocal microscopy at an excitation/emission wavelength of 494/506 nm.

### 2.12 Quantitative filamentous actin/globular actin (F-actin/G-actin) ratio measurement

The F-actin/G-actin ratio was measured using G-Actin/F-Actin In Vivo Assay Biochem Kit following the manufacturer’s instructions.

### 2.13 Statistical analysis

The data are reported as the mean ± standard error of the mean (SEM) from a minimum of three to five separate experiments. Graphs were produced using GraphPad Prism 8. Statistical significance was determined through either paired or unpaired Student’s t-tests, with a threshold of p ≤ 0.05 indicating significance.

**Project accession:** MassIVE MSV000096855 Username: MSV000096855_reviewer Password: CTSK

## 3. Results

### 3.1. Differential expression of proteins in TM subjected to functional loss of CTSK

Tandem mass tag (TMT)-based unbiased global proteomic analysis of HTM cells treated with siCTSK identified 5,949 proteins, of which 5,843 were quantified. Based on FDR ≤5%, statistical significance (p ≤ 0.050), and a confidence threshold of log2 fold change (log2FC) outside the mean ± 2σ (log2FC ≤ -0.1 or ≥ 0.1), 324 proteins were significantly upregulated and 82 were significantly downregulated in TM under siCTSK treatment compared to siScr. **Table 2** shows the top 20 upregulated proteins.

**Table 2:**
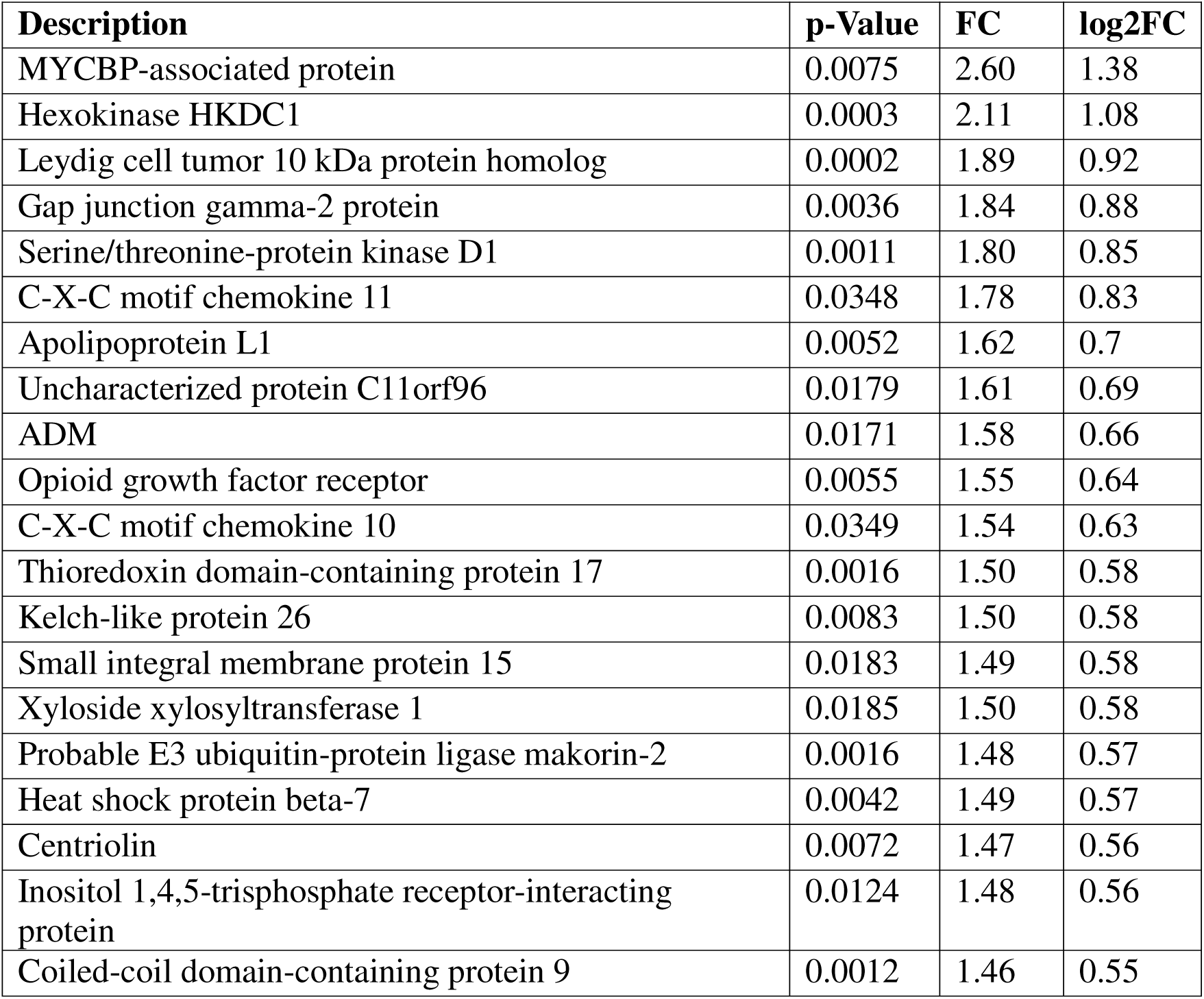
Top 20 upregulated proteins.

| **Description** | **p-Value** | **FC** | **log2FC** |
| --- | --- | --- | --- |
| MYCBP-associated protein | 0.0075 | 2.60 | 1.38 |
| Hexokinase HKDC1 | 0.0003 | 2.11 | 1.08 |
| Leydig cell tumor 10 kDa protein homolog | 0.0002 | 1.89 | 0.92 |
| Gap junction gamma-2 protein | 0.0036 | 1.84 | 0.88 |
| Serine/threonine-protein kinase D1 | 0.0011 | 1.80 | 0.85 |
| C-X-C motif chemokine 11 | 0.0348 | 1.78 | 0.83 |
| Apolipoprotein L1 | 0.0052 | 1.62 | 0.7 |
| Uncharacterized protein C11orf96 | 0.0179 | 1.61 | 0.69 |
| ADM | 0.0171 | 1.58 | 0.66 |
| Opioid growth factor receptor | 0.0055 | 1.55 | 0.64 |
| C-X-C motif chemokine 10 | 0.0349 | 1.54 | 0.63 |
| Thioredoxin domain-containing protein 17 | 0.0016 | 1.50 | 0.58 |
| Kelch-like protein 26 | 0.0083 | 1.50 | 0.58 |
| Small integral membrane protein 15 | 0.0183 | 1.49 | 0.58 |
| Xyloside xylosyltransferase 1 | 0.0185 | 1.50 | 0.58 |
| Probable E3 ubiquitin-protein ligase makorin-2 | 0.0016 | 1.48 | 0.57 |
| Heat shock protein beta-7 | 0.0042 | 1.49 | 0.57 |
| Centriolin | 0.0072 | 1.47 | 0.56 |
| Inositol 1,4,5-trisphosphate receptor-interacting protein | 0.0124 | 1.48 | 0.56 |
| Coiled-coil domain-containing protein 9 | 0.0012 | 1.46 | 0.55 |

Under siCTSK treatment, the most significantly upregulated protein was MYCBP-associated protein (log2FC: 1.38; FC: 2.6). MYCBPAP is primarily known for its role in spermatogenesis and is also predicted to contribute to transcriptional regulation through its interaction with MYC-binding protein (16). The second most upregulated protein was Hexokinase HKDC1 (log2FC: 1.08; FC: 2.11) which is involved in glucose homeostasis. Its upregulation has been reported to contribute to epithelial-mesenchymal transition (EMT) and fibrosis in several cancers, including liver cancer, gastric cancer and lung adenocarcinoma (17–19). This was followed by Leydig cell tumor 10 kDa protein homolog (log2FC: 0.92; FC: 1.89), a protein implicated in steroid biosynthesis and testicular function (20). While its precise role outside reproductive tissues remains unclear, it has been associated with intracellular signaling and may contribute to cellular metabolism and stress response (21).

Among the top upregulated proteins was serine/threonine-protein kinase D1 (PRKD1) (log2FC: 0.85; FC: 1.80). PRKD1 is a kinase that phosphorylates various proteins and regulates a variety of cellular functions, including membrane receptor signaling, protection from oxidative stress, gene transcription, and cytoskeletal dynamics (22, 23). In addition to PRKD1, several other actin-cytoskeleton-associated proteins were upregulated, including WAS/WASL-interacting protein family member 2 (WIPF2) (log2FC: 0.21; FC: 1.16), which is involved in actin polymerization (24); paxillin (PXN) (log2FC: 0.19; FC: 1.14), a focal adhesion adaptor protein; FYVE, RhoGEF, and PH domain-containing protein 4 (FGD4) (log2FC: 0.17; FC: 1.12), a guanine nucleotide exchange factor involved in cytoskeleton rearrangement (25); phosphatidylinositol 4-phosphate 5-kinase type-1 alpha (PIP5K1A) (log2FC: 0.15; FC: 1.11), which is involved in actin remodeling by synthesizing phosphatidylinositol 4,5-bisphosphate (PIP2) (26); tensin-1 (TNS1) (log2FC: 0.12; FC: 1.09), which plays a key role in fibrillar adhesion formation by linking the actin cytoskeleton to extracellular fibronectin fibrils (27); phostensin (PPP1R18) (log2FC: 0.12; FC: 1.09), an actin-binding regulatory subunit of protein phosphatase 1 (28); and [F-actin]-monooxygenase MICAL1 (log2FC: 0.11; FC: 1.08), an oxidoreductase that controls actin disassembly (29).

ECM associated proteins were also enriched in the upregulated dataset, including SPARC-related modular calcium-binding protein 2 (SMOC2) (log2FC: 0.4; FC: 1.32), an ECM-associated, calcium-binding protein involved in collagen biogenesis. SMOC2 potentially induces collagen maturation by modulating bone morphogenetic protein 1 (BMP1) activity and the biogenesis of other ECM proteins. Polymorphisms in the SMOC2 gene have been associated with glaucoma (30, 31). Also upregulated was lysyl oxidase homolog 3 (LOXL3) (log2FC: 0.32; FC: 1.25), an enzyme involved in collagen cross-linking (32). Epidermal growth factor-like protein 7 (EGFL7) (log2FC: 0.26; FC: 1.20), which plays a crucial role in modulating interactions with the ECM, was also enriched. Additionally, mothers against decapentaplegic homolog 3 (SMAD3) (log2FC: 0.12; FC: 1.09), which is a key regulator of transforming growth factor β2 (TGFβ2) signaling, was upregulated. Notably, TGFβ2-induced IOP elevation has been shown to be dependent on SMAD3 (33, 34), was upregulated. Finally, SH3 and PX domain-containing protein 2B (SH3PXD2B) (log2FC: 0.11; FC: 1.08), an adaptor protein associated with matrix remodeling (35), was significantly increased.

Proteins related to cell adhesion were also among the upregulated targets. Intercellular adhesion molecule 1 (ICAM1) (log2FC: 0.43; FC: 1.35) and intercellular adhesion molecule 5 (ICAM5) (log2FC: 0.26; FC: 1.20), both known to mediate cell-cell interactions, were increased, along with nectin-2 (log2FC: 0.13; FC: 1.10), which plays a role in cell adhesion through nectin signaling (36), and cytoplasmic protein NCK1 (log2FC: 0.1; FC: 1.07), an adaptor protein that links cell adhesion complexes to actin cytoskeleton remodeling (37).

Interestingly, in the context of immune regulation, several proteins known to influence inflammatory responses were elevated. Macrophage migration inhibitory factor (MIF) (log2FC: 0.34; FC: 1.27), a pro-inflammatory cytokine involved in immune cell activation (38), and macrophage colony-stimulating factor 1 (CSF1) (log2FC: 0.3; FC: 1.23), a cytokine critical for macrophage differentiation (39), were significantly increased.

Additionally, C-X-C motif chemokine 11 (CXCL11) (log2FC: 0.63; FC: 1.54) and C-X-C motif chemokine 10 (CXCL10) (log2FC: 0.34; FC: 1.27), which primarily signal through CXCR3 and, to a lesser extent, ACKR3, play key roles in immune cell recruitment and inflammatory responses. While their main functions involve immune regulation, some studies suggest that they may also influence intracellular calcium mobilization (40, 41). In addition to these, In addition, inositol 1,4,5-trisphosphate receptor-interacting protein (ITPRIP) (log2FC: 0.56; FC: 1.48), which plays a crucial role in regulating calcium release from the endoplasmic reticulum by interacting with the inositol 1,4,5-trisphosphate receptor (IP3R) (42), was also significantly upregulated.

Furthermore, adrenomedullin (ADM) (log2FC: 0.66; FC: 1.58), a multifunctional peptide involved in vasodilation, oxidative stress protection, and calcium homeostasis (43, 44), was significantly upregulated in siCTSK-treated TM cells. ADM has been reported to modulate endothelial barrier function and inflammatory signaling, and its increase may reflect a stress response to ECM remodeling and calcium dysregulation (45, 46). Given its role in regulating cytoskeletal integrity and cell survival, ADM upregulation may contribute to the adaptive changes observed following CTSK depletion.

Finally, proteins associated with apoptosis were notably upregulated, suggesting potential changes in cell survival mechanisms under siCTSK treatment. These included caspase-7 (CASP7) (log2FC: 0.28; FC: 1.21), an executioner caspase involved in apoptosis (47); tumor necrosis factor receptor type 1-associated DEATH domain protein (TRADD) (log2FC: 0.16; FC: 1.12), a key adaptor in TNF-induced apoptosis (48); protein phosphatase 1F (PPM1F) (log2FC: 0.15; FC: 1.11), a regulator of stress-induced apoptotic signaling (49); caspase-3 (CASP3) (log2FC: 0.12; FC: 1.09), another executioner caspase involved in apoptosis (47); and apoptosis inhibitor 5 (API5) (log2FC: 0.12; FC: 1.09), which negatively regulates apoptosis (50).

Pathway enrichment analysis using ShinyGO (51) for upregulated proteins in TM induced by siCTSK treatment based on molecular function is given in **Table 3**. The enriched pathways predominantly include proteins associated with nucleic acid binding, enzyme binding, carbohydrate derivative binding, and ATP binding. Additionally, several proteins involved in cytoskeletal protein binding, kinase activity, and transferase activity related to phosphorus-containing groups were enriched. Other significantly enriched functions include protein domain-specific binding, GTPase binding, heat shock protein binding, and unfolded protein binding, indicating potential roles in cellular stress response and protein homeostasis. Furthermore, fibronectin binding was also enriched, highlighting potential ECM interactions and signaling pathways. Notably, enrichment of snRNA binding and translation initiation factor binding suggests potential regulation at the RNA-processing and translational level.

**Table 3:**
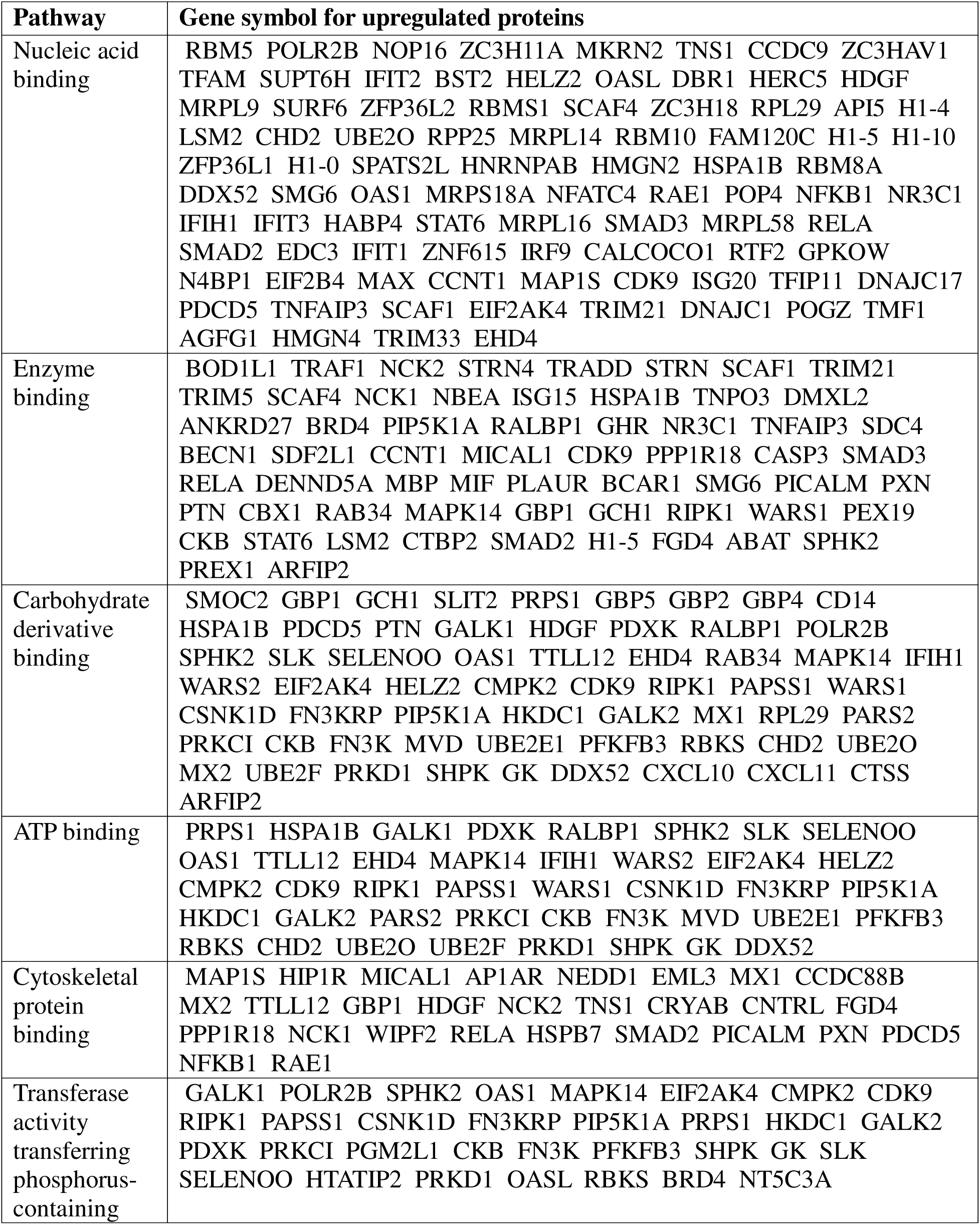
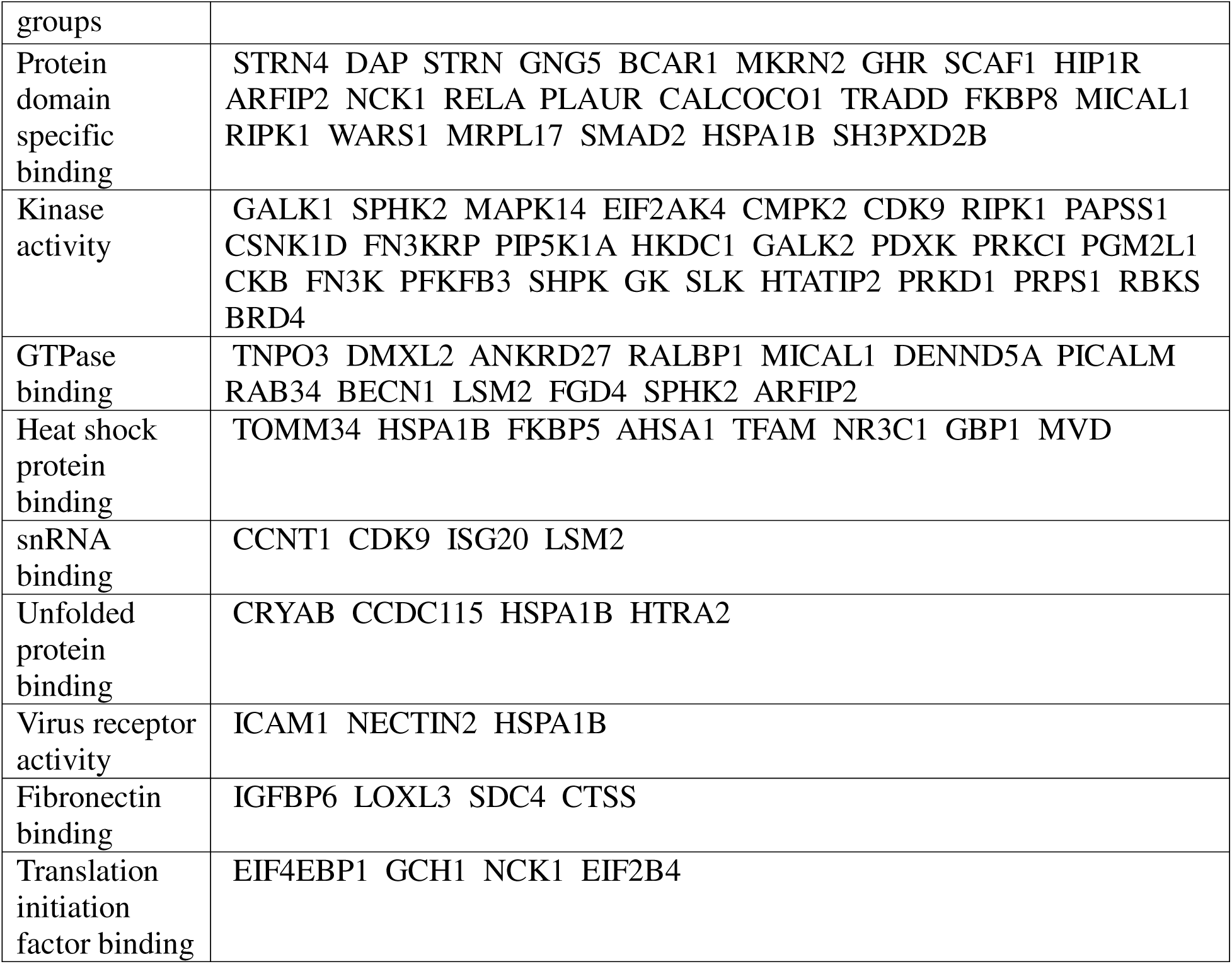
Pathway enrichment analysis for upregulated proteins using ShinyGO based on molecular function.

On examining the 82 downregulated proteins, we found that CTSK (log2FC: -0.61; FC: 0.66), the primary target of siCTSK treatment, was among the most significantly reduced proteins **(Table 4)**. In addition to CTSK, several other actin-cytoskeleton-associated proteins were downregulated, including Myosin-7 (log2FC: -1.59; FC: 0.33) and Myosin-6 (log2FC: - 0.91; FC: 0.53) were the two most significantly downregulated proteins. These molecular motors interact with actin filaments and play critical roles in cytoskeletal organization, intracellular transport, and cellular contractility (52, 53). Myosin-13 (MYH13) (log2FC: -0.45; FC: 0.73) and Rho-related GTP-binding protein RhoQ (RHOQ) (log2FC: -0.37; FC: 0.78), a small GTPase involved in actin cytoskeletal reorganization, were also significantly reduced. Alpha-catulin (CTNNAL1) (log2FC: -0.15; FC: 0.90), an actin-associated protein linked to cell motility and signaling (54), was also significantly downregulated.

**Table 4:**
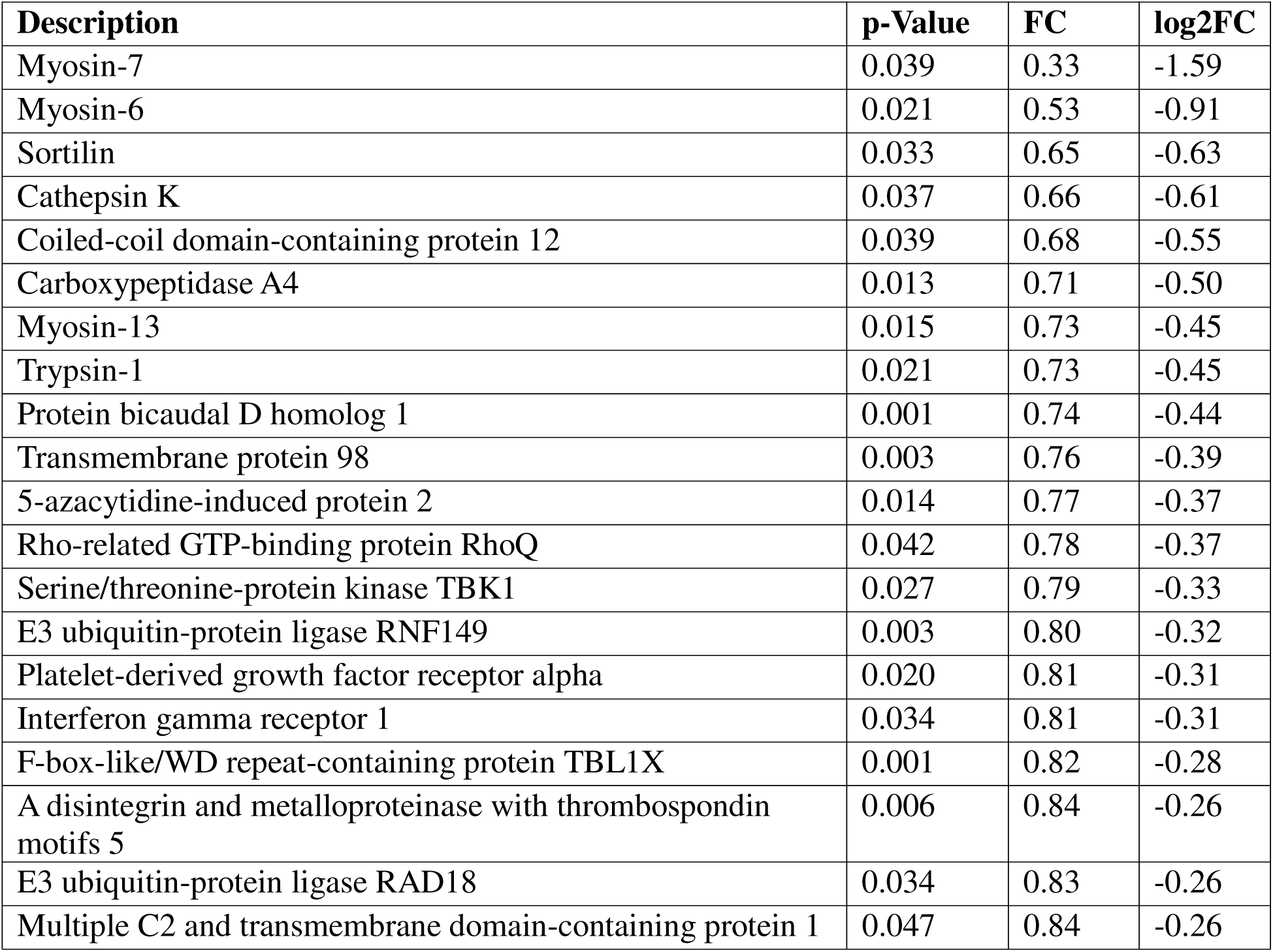
Top 20 downregulated proteins.

A significant proportion of the downregulated proteins were associated with the ECM, including a disintegrin and metalloproteinase with thrombospondin motifs 5 (ADAMTS5) (log2FC: -0.26; FC: 0.84), an ECM protease involved in proteoglycan degradation (55) and, collagen alpha-1(VI) chain (COL6A1) (log2FC: -0.23; FC: 0.85), a key structural component of the ECM that supports tissue integrity, were significantly reduced. Additionally, significant downregulation of prolyl 4-hydroxylase subunit alpha-2 (P4HA2) (log2FC: -0.17; FC: 0.89), an enzyme critical for stability of collagen (56), was observed. Similarly, reversion-inducing cysteine-rich protein with Kazal motifs (RECK) (log2FC: -0.16; FC: 0.90), a metalloproteinase inhibitor involved in ECM homeostasis (57), and disintegrin and metalloproteinase domain-containing protein 9 (ADAM9) (log2FC: -0.16; FC: 0.89), which functions in ECM remodeling and proteolysis (58), were also significantly downregulated. Furthermore, procollagen C-endopeptidase enhancer 1 (PCOLCE) (log2FC: -0.16; FC: 0.90), which regulates collagen maturation (59), was also reduced significantly. Consistently, PCOLCE1 downregulation has also been observed in TM cells treated with TGFβ2 (60).

Beyond ECM remodeling, two proteins involved in cell-ECM interactions were also downregulated-tetraspanin-6 (TSPAN6) (log2FC: -0.18; FC: 0.89), which plays a role in cell adhesion and membrane trafficking (61), and integrin beta-1 (ITGB1) (log2FC: -0.10; FC: 0.93), a key receptor facilitating cell-ECM interactions (62).

Additionally, several signaling, and regulatory proteins were downregulated. Serine/threonine-protein kinase TBK1 (log2FC: -0.33; FC: 0.79), a key player in immune and stress signaling pathways (63), was downregulated. Platelet-derived growth factor receptor alpha (PDGFRA) (log2FC: -0.31; FC: 0.81), which regulates cell proliferation and migration (64), was also significantly downregulated. Similarly, E3 ubiquitin-protein ligase RAD18 (log2FC: -0.23; FC: 0.86), which plays a role in DNA damage repair and post-replication repair, and E3 ubiquitin-protein ligase CBL (log2FC: -0.19; FC: 0.88), involved in protein degradation and receptor signaling, were also downregulated.

Pathway enrichment analysis using ShinyGO for downregulated proteins in TM induced by siCTSK treatment based on molecular function is given in **Table 5**. Among the enriched pathways are proteins associated with protein-containing complex binding, collagen binding, and growth factor binding, indicating a reduced capacity for structural and signaling interactions within the TM. The enrichment of metallopeptidase activity and ECM binding suggests a reduced ECM remodeling capacity. The significant enrichment of microfilament motor activity suggests a potential impairment in actin cytoskeletal dynamics and intracellular transport mechanisms.

**Table 5:** Pathway enrichment analysis for downregulated proteins using ShinyGO based on molecular function.

| **Pathway** | **Gene symbol for downregulated proteins** |
| --- | --- |
| Protein-containing complex binding | PCOLCE ITGB1 BICD1 ADAM9 SORT1 CTSK PI4K2A MYH13 MYH7 CTNNAL1 CLTA PDGFRA PRNP MYH6 RAD18 COL6A1 ADAMTS5 |
| Collagen binding | PCOLCE ITGB1 CTSK ADAM9 COL6A1 |
| Growth factor binding | IL6ST PDGFRA COL6A1 A2M SORT1 |
| Metallopeptidase activity | MIPEP CPD CPA4 ADAMTS5 ADAM9 |
| Microfilament motor activity | MYH7 MYH6 MYH13 |
| Extracellular matrix binding | ITGB1 ADAMTS5 ADAM9 |

### 3.2 Functional loss of CTSK disrupted collagen biogenesis and ECM homeostasis

Our previous study showed increase in ECM proteins under CTSK activity inhibition and reduction in ECM proteins when CTSK was induced (9). Moreover, CTSK is a potent collagenase. Hence, we wanted to analyze the effect of functional loss of CTSK on collagen biogenesis.

To validate the efficiency of CTSK knockdown, we examined its expression in both whole cell lysate (WCL) and conditioned media (CM) via immunoblot (IB) analysis. A significant reduction in both the pro (p=0.002) and active (p=0.01) forms of CTSK was observed in siCTSK-treated TM cells compared to siScr control **(Figure 1A)**. The reduction of pro-CTSK (p=0.01) in CM further validates the knockdown efficiency at the secretory level, which is particularly relevant given the role of CTSK in ECM turnover **(Figure 1B)**.

**Figure 1.**
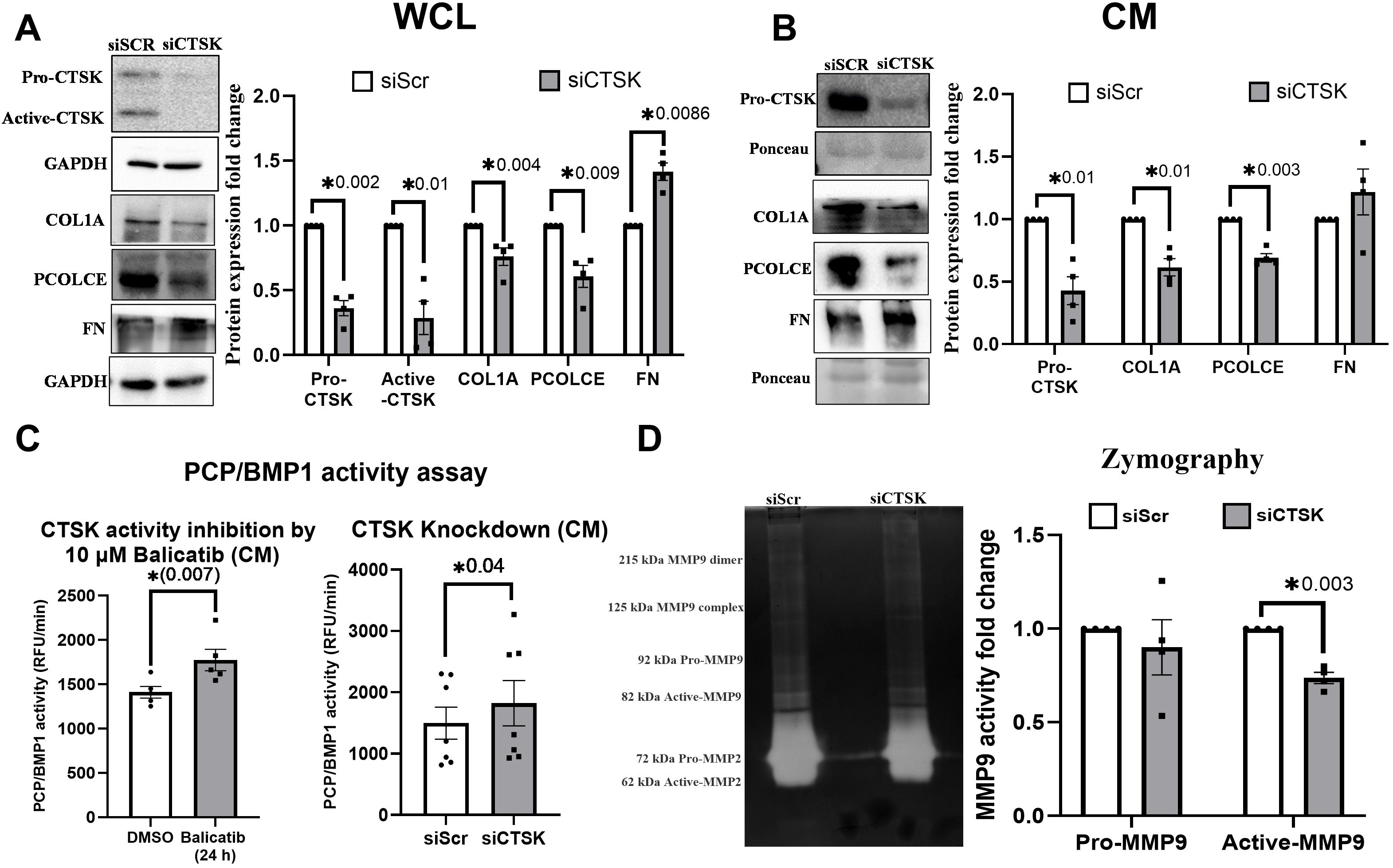
Functional loss of CTSK disrupts collagen biogenesis and ECM homeostasis. (A) Immunoblot (IB) analysis of whole-cell lysate (WCL) showing a significant reduction in pro-CTSK and active-CTSK upon siCTSK treatment compared to siScr control. Collagen I (COL1A) and PCOLCE levels were significantly downregulated, whereas Fibronectin (FN) was upregulated in siCTSK-treated cells. GAPDH was used as a loading control. (B) IB analysis of conditioned media (CM) showing a reduction in secreted pro-CTSK, COL1A, and PCOLCE following CTSK knockdown. (C) PCP/BMP1 activity assay performed using conditioned media from Balicatib (10 μM), siScr- and siCTSK-treated cells. BMP1 activity (RFU/min) was significantly increased upon CTSK activity inhibition and knockdown. (D) Zymography analysis of conditioned media showing a significant reduction in active-MMP9 levels in siCTSK-treated cells compared to siScr control, indicating a potential role for CTSK in MMP9 activation and ECM remodeling. Data are presented as mean ± SEM. Statistical significance was achieved using a paired Student’s t-test, with a sample size of 4–7 per experiment.

CTSK knockdown resulted in significant changes in collagen biogenesis and ECM composition. Collagen I (COL1A) (p=0.004) and PCOLCE (p=0.009) were significantly downregulated in WCL, suggesting a disruption in collagen synthesis or maturation **(Figure 1A)**. Conversely, fibronectin (FN) (p=0.0086) was significantly upregulated in siCTSK.

To assess whether the reduction in COL1A and PCOLCE also affects their secretion, IB was performed on conditioned media **(Figure 1B)**. Consistent with WCL findings, both COL1A (p=0.01) and PCOLCE (p=0.003) were significantly reduced in CM, suggesting an overall reduction in extracellular collagen deposition and processing. Interestingly, FN levels were increased in CM, but this change was not statistically significant. The decrease in secretory COL1A and PCOLCE implies that collagen maturation is disrupted, potentially contributing to ECM dysregulation following CTSK depletion.

To further investigate the effect of CTSK loss on collagen biogenesis, we assessed the activity of bone morphogenetic protein 1 (BMP1), a crucial enzyme in collagen maturation (65) **(Figure 1C)**. Given that BMP1 plays a key role in collagen processing, and our previous study showed an increase in collagen expression when CTSK activity was pharmacologically inhibited using Balicatib, we hypothesized that BMP1 activity might be altered in siCTSK-treated cells. We performed a PCP/BMP1 activity assay using CM, which serves as the most relevant sample for assessing extracellular collagen-processing enzymes. BMP1 activity was significantly increased under both CTSK activity inhibition with 10 μM Balicatib (p=0.007) for 24h and siRNA-mediated CTSK knockdown (p=0.04) conditions.

Matrix metalloproteinases (MMPs) play a key role in ECM turnover and degradation (66), and we sought to determine whether CTSK loss affects MMP activity. Zymography analysis of conditioned media **(Figure 1D)** demonstrated a significant reduction in active-MMP9 levels (p=0.003) in siCTSK-treated cells compared to siScr, suggests that CTSK plays a crucial role in MMP9 activation, a finding that aligns with previous studies (67).

### 3.3 CTSK depletion results in sustained elevation in intracellular calcium levels in TM cells

Changes in ECM composition can alter integrin-mediated signaling and mechanotransduction, which are known to impact intracellular calcium homeostasis (68). Given the ECM alterations observed upon CTSK knockdown, we next investigated whether intracellular calcium levels were affected.

Relative fluorescence unit (RFU) measurements confirmed that acetylcholine chloride (5 μM), used as a positive control, significantly increased intracellular calcium levels (p < 0.05) compared to non-treated cells **(Figure 2A)**. Notably, both CTSK activity inhibition with balicatib (10 μM, p < 0.05; **Figure 2B**) and siCTSK treatment for 12, 24, and 48h (p < 0.05; **Figure 2C**) led to a significant increase in intracellular calcium levels.

**Figure 2.**
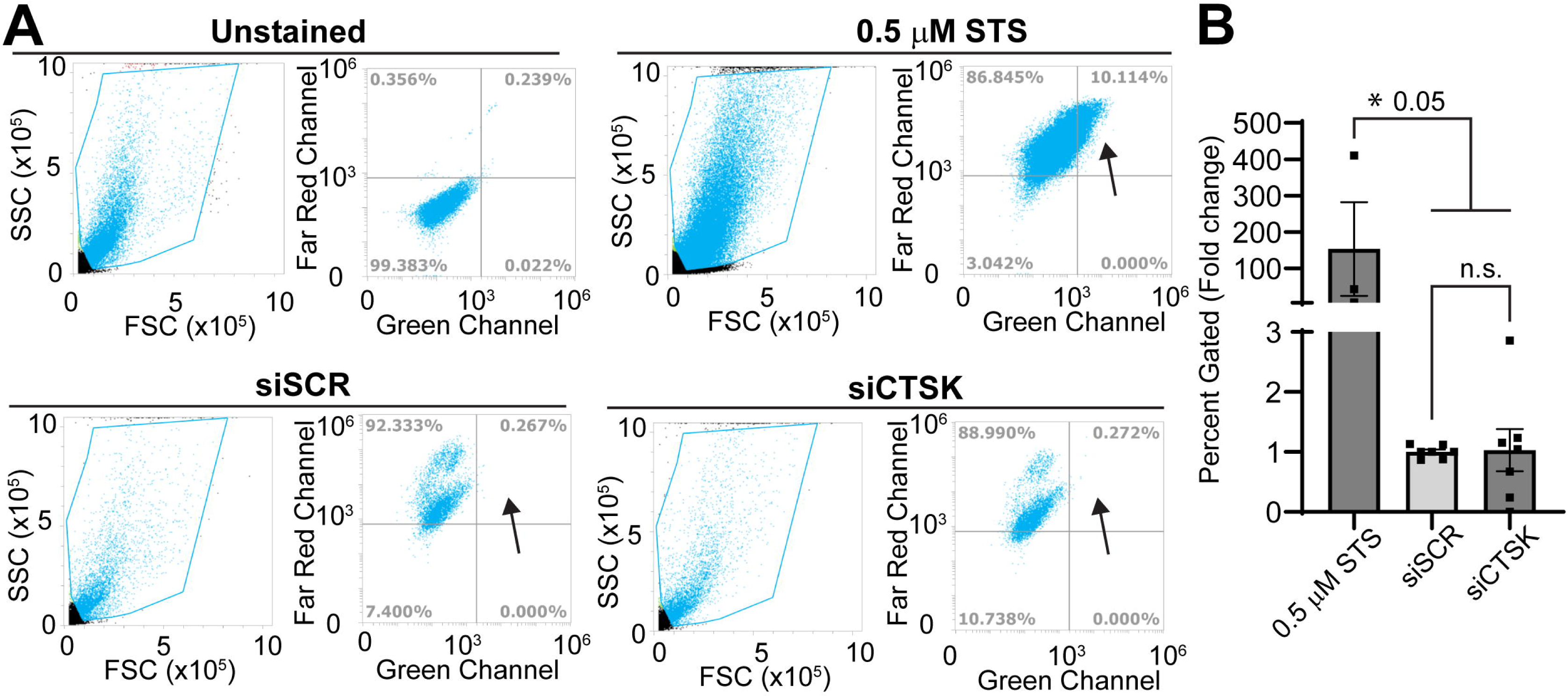
CTSK depletion elevates intracellular calcium levels in TM cells. (A) Relative fluorescence unit (RFU) measurements using a microplate reader confirmed that acetylcholine (5 μM, positive control) significantly increased intracellular calcium levels compared to non-treated cells. (B) CTSK activity inhibition using Balicatib (10 μM) significantly increased intracellular calcium levels compared to DMSO control. (C) siCTSK-treated TM cells at 12, 24, and 48h exhibited a significant increase in intracellular calcium levels compared to siScr controls at respective time points. The violin plot displays median fluorescence intensity (red line). (D) Confocal imaging of intracellular calcium levels, showing increased Fluo-4 AM fluorescence (green) intensity in acetylcholine- and Balicatib-treated cells compared to non-treated and DMSO controls, respectively. Yellow arrows indicate regions of enhanced fluorescence. (E) Confocal imaging of siCTSK-treated cells (24h and 48h) showing increased intracellular calcium fluorescence intensity compared to siScr-treated cells at the same time points. Data are represented as mean ± SEM, with statistical significance achieved through unpaired student’s t test with sample size of n = 5-9.

Confocal imaging further validated these findings, showing a marked increase in green fluorescence intensity indicative of elevated intracellular calcium levels. **Figure 2D** highlights enhanced fluorescence upon acetylcholine treatment and Balicatib treatment compared to non-treated and DMSO controls, respectively. Similarly, **Figure 2E** demonstrates increased intracellular calcium levels in siCTSK-treated cells at both 24 and 48h compared to siScr controls at the same time points.

These results suggest that CTSK depletion disrupts calcium homeostasis in TM cells, leading to sustained elevation in intracellular calcium levels, which could have downstream effects on cellular signaling and mechanotransduction. While calcium signaling can be rapidly induced, our approach focused on intracellular calcium changes at 12-, 24-, and 48h post-siRNA transfection to assess sustained effects rather than transient fluctuations immediately following CTSK depletion.

### 3.4 CTSK loss induces actin polymerization through PRKD1 activation

Since intracellular calcium plays a critical role in cytoskeletal dynamics, we next investigated whether CTSK depletion induces actin polymerization and its regulatory pathways. Proteomic analysis showed a significant increase in PRKD1 in siCTSK-treated cells. To confirm its activation, we analyzed the phosphorylation status of PRKD1 (p-PRKD1; S744/748). Both PRKD1 (p=0.04) and p-PRKD1 (p=0.03) showed a significant increase in siCTSK compared to siScr **(Figure 3A)**. However, the p-PRKD1/PRKD1 ratio did not show a significant change, suggesting that while total PRKD1 levels increased, the proportion of active PRKD1 remained relatively stable. Nonetheless, the observed increase in p-PRKD1 levels supports its activation and potential downstream effects.

**Figure 3.**
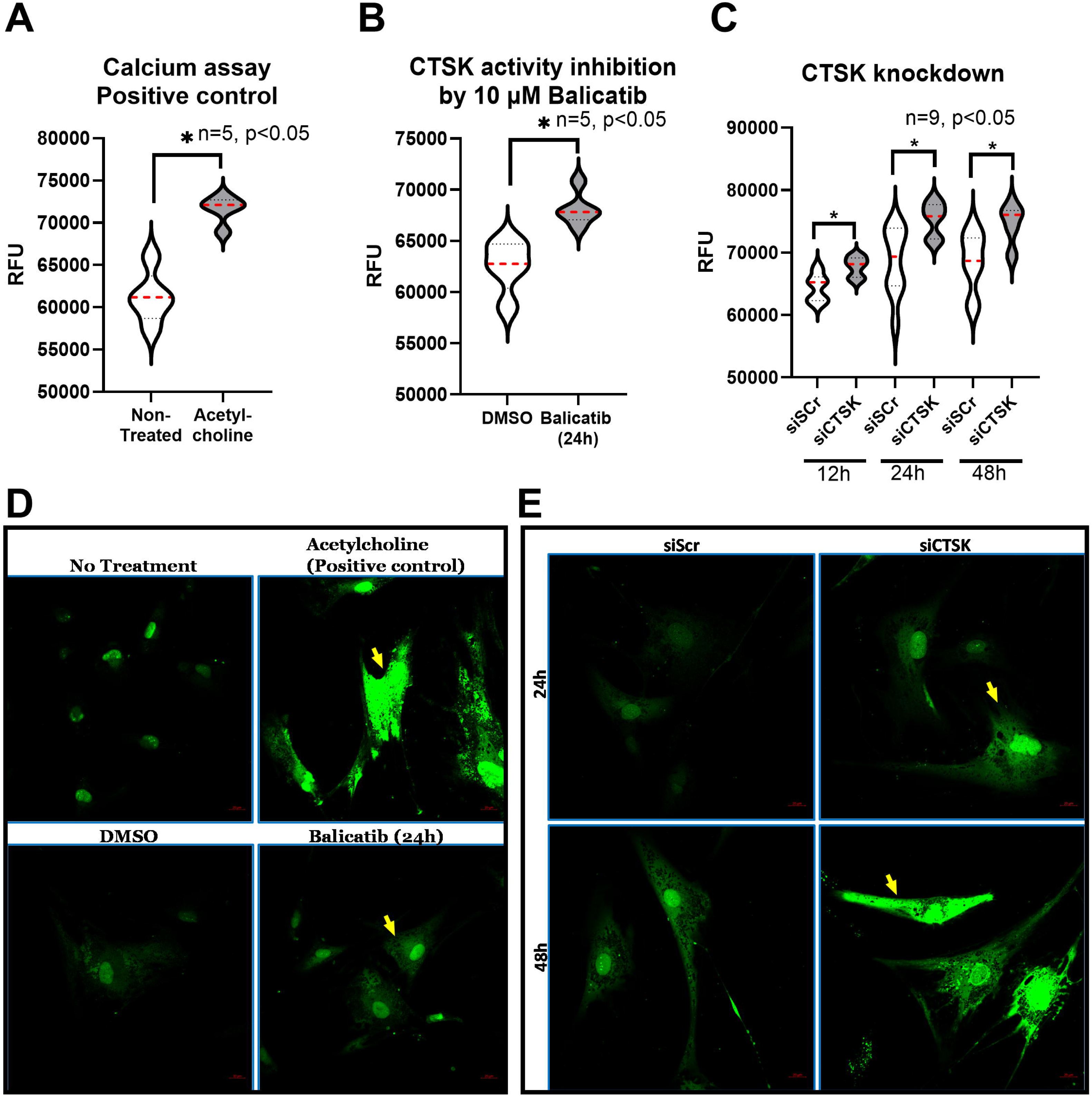
CTSK loss induces actin polymerization through PRKD1 activation. (A) Immunoblot (IB) analysis of PRKD1 and its downstream targets in siScr- and siCTSK-treated TM cells. Significant increase in i) PRKD1, ii) p-PRKD1, iii) p-SSH1, iv) p-SSH1/SSH1 ratio, v) p-cofilin, and vi) p-cofilin/cofilin ratio were observed in siCTSK treatment compared to siScr. GAPDH was used as a loading control. (B) Immunofluorescence (IF) staining for CTSK (red) and actin filaments (phalloidin, green). Phalloidin fluorescence intensity increased in siCTSK-treated cells (red arrows), suggesting enhanced actin polymerization. Grayscale images are included for clarity. Quantification using CellProfiler v4.2.7 confirmed a significant increase. (C) F-actin/G-actin assay showing an increase in F-actin levels and F-actin/G-actin ratio in siCTSK-treated cells. GAPDH was used as a loading control but not for quantification. (D) IF staining of vinculin (red) and actin filaments (phalloidin, green) in siScr- and siCTSK-treated TM cells. Increased vinculin puncta (red arrows) were observed in siCTSK-treated cells, especially at focal adhesions. Vinculin puncta were quantified using CellProfiler (Otsu threshold method), showing a significant increase in siCTSK-treated cells. Statistical analysis: Data are presented as mean ± SEM, with p < 0.05 considered significant (n = 3–5 biological replicates). Paired Student’s t-test was used for IB analysis, while unpaired Student’s t-test was used for IF quantification.

PRKD1 has been demonstrated to exert substantial control over a wide range of cellular functions including intracellular signaling, proliferation, and invasive behavior (69) and play an important role in actin reorganization (22). Actin remodeling involves phosphorylation of LIM kinase 1 (LIMK1) to promote actin polymerization and slingshot phosphatase 1 (SSH1) suppressing cofilin, an actin-severing protein (70). To investigate these downstream effects, we examined the protein expression and phosphorylation status of LIMK1, SSH1, and cofilin **(Figure 3A)**. While both LIMK1 and p-LIMK1 (Thr508) were elevated in siCTSK-treated cells, these changes were not statistically significant, possibly due to temporal variations in cellular signaling. In contrast, p-SSH1 (Ser978) (p = 0.05) was significantly elevated, leading to an increase in the p-SSH1/SSH1 ratio (p = 0.02) in siCTSK-treated cells. Although total cofilin levels remained unchanged, p-cofilin (Ser3) (p = 0.01) was significantly increased in siCTSK-treated cells, and the p-cofilin/cofilin ratio (p = 0.002) was significantly elevated compared to siScr (**Figure 3A**). These findings indicate that CTSK depletion can result in increased PRKD1 activity, which may contribute to enhanced actin polymerization through the LIMK1/SSH1/cofilin pathway.

To visualize actin cytoskeletal changes, immunofluorescence staining was performed for CTSK (red) and actin filaments (phalloidin, green) **(Figure 3B)**. Representative images show siScr-treated cells in the top panel and siCTSK-treated cells in the bottom panel. Phalloidin fluorescence intensity was markedly increased in siCTSK-treated cells (indicated by red arrows) compared to siScr controls. Grayscale images of phalloidin staining are included for clarity. Fluorescence intensity of actin fibers was quantified using CellProfiler v4.2.7, which showed a significant increase (p = 0.02) in siCTSK-treated cells (graph on the far right).

To further assess actin polymerization, we measured the F-actin/G-actin ratio **(Figure 3C)**. F-actin intensity was increased in siCTSK-treated cells, and the F-actin/G-actin ratio (p = 0.03) was significantly elevated compared to siScr controls. Experimental positive control blots (not shown) confirmed effective separation of F-actin and G-actin fractions. GAPDH from the G-actin blot was used to show equal loading but was not used for relative quantification.

Given the increase in actin polymerization and previous findings on changes in ECM remodeling in the absence of CTSK, we next investigated the distribution of the focal adhesion protein vinculin (71), which plays a key role in focal adhesion dynamics and cytoskeletal organization. Immunofluorescence analysis showed an increase in vinculin puncta (red) along phalloidin-stained actin filaments (green) in siCTSK-treated cells **(Figure 3D)**. The top panel shows siScr-treated cells, while the bottom panel shows siCTSK-treated cells. Grayscale images of phalloidin and vinculin staining are provided for clarity. Red arrows indicate vinculin puncta. Vinculin puncta were quantified using CellProfiler v4.2.7 with the Otsu threshold method, and the graph on the far right shows a significant increase in vinculin puncta count (p = 0.05) in siCTSK-treated cells compared to siScr controls.

These findings reinforced that CTSK depletion enhances actin polymerization and focal adhesion formation, potentially through PRKD1 activation and its downstream effects on LIMK1, SSH1, and cofilin.

### 3.5 CTSK loss increases apoptotic markers but does not induce apoptosis in TM cells

PRKD1 activation and actin remodeling are known to regulate apoptotic pathways, and our proteomic analysis showed an increase in apoptotic markers such as caspases (CASP3 and CASP7), TRADD, and PPM1F, we next examined whether CTSK loss induced apoptosis in TM cells. This was carried out using flow cytometry analysis of caspase 3/7 activation in siCTSK-, siSCR-, and 0.5 μM staurosporine (STS, positive control)-treated HTM cells. Cells were stained with NucRed (nuclear stain) and CellEvent Caspase 3/7 Green detection reagent. Cells were gated to exclude debris, and unstained controls were used to define the population positive for both NucRed and caspase 3/7 (**Figure 4A**, indicated by arrows). Our analysis revealed no significant increase in the percentage of apoptotic (NucRed^+^ and Caspase^+^) cells in siCTSK-treated cells compared to siSCR controls **(Figure 4A)**. STS treatment, as expected, induced a significant increase in caspase 3/7 activation (p = 0.05) compared to both siScr and siCTSK-treated cells, whereas siCTSK treatment did not show a similar effect. Quantification of the double-positive apoptotic cell population **(Figure 4B)**, normalized to siSCR controls, confirmed that siCTSK does not induce apoptosis in HTM cells.

**Figure 4.**
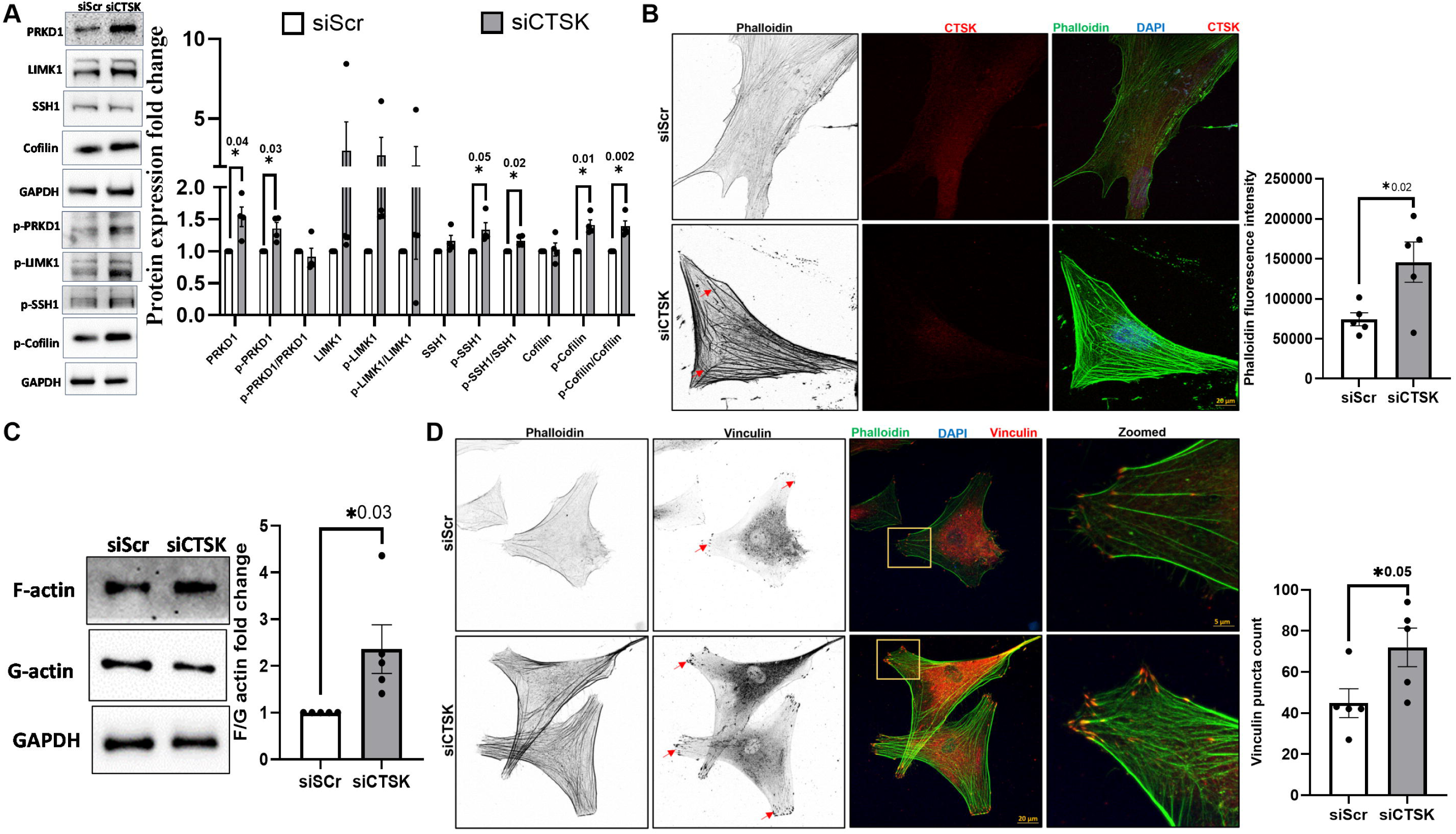
CTSK loss increases apoptotic markers but does not induce apoptosis in TM cells. (A) HTM cells were stained with NucRed (Far Red) and Caspase Detection Reagent (Green). Cell debris was gated out first and then unstained cells were used to create NucRed+/Caspase+ gates (indicated by the arrows) for the STS, siSCR, and siCTSK treated cells. (B) The percentage of cells gated into the NucRed+/Caspase+ population, relative to that of siSCR, show siCTSK does not increase apoptosis, whereas STS treatment induced a significant increase (p < 0.05). One-way Anova with Tukey’s, n= 3-7, * p < 0.05. FSC-Forward Scatter; SSC-Side Scatter.

In summary, our findings indicate that while CTSK loss increases apoptotic markers, it does not induce apoptosis in TM cells under the conditions tested. These results suggest that the ECM and cytoskeletal changes observed in CTSK knockdown cells occur independently of apoptotic cell death. Further investigation is needed to determine the mechanisms underlying apoptotic marker upregulation and the broader functional consequences of CTSK depletion in HTM cells.

## 4. Discussion

In this study, we sought to determine the molecular consequences of CTSK loss in TM cells using global proteomics analysis, biochemical assays, and functional validation. Our findings demonstrate that CTSK depletion alters actin cytoskeletal dynamics, ECM remodeling, intracellular calcium homeostasis, and focal adhesion assembly.

CTSK is a potent collagenase, and in our previous study, we demonstrated that inhibition of CTSK activity increases collagen deposition and actin bundling and induction of CTSK reduces ECM accumulation (9). In this study, we show that functional loss of CTSK significantly impacted collagen processing and ECM remodeling. We observed a significant reduction in COL1A and PCOLCE protein levels in both WCL and CM, suggesting impaired collagen synthesis and secretion. PCOLCE is a key modulator of collagen fibrillogenesis (59), and its reduction may contribute to ECM destabilization. In contrast, fibronectin was significantly upregulated, which may represent a compensatory response to ECM disruption. To further investigate collagen-processing mechanisms, we assessed BMP1 activity, which is crucial for collagen maturation. BMP1 activity was significantly elevated in both pharmacological CTSK inhibition (balicatib) and siRNA-mediated CTSK knockdown. These findings suggest that BMP1 activation may act as a compensatory mechanism to mitigate the effects of reduced PCOLCE expression and impaired collagen maturation. It is possible that other cathepsins or ECM-degrading proteases may also contribute to collagen remodeling in the absence of CTSK. Additionally, our proteomics analysis revealed an upregulation of ECM-associated proteins SMOC2 and LOXL3, which may reflect an adaptive response to maintain ECM homeostasis under CTSK depletion. LOXL3 plays a role in collagen cross-linking (32, 72), while SMOC2 antagonizes BMP1 signaling (73). Increased levels of LOXL3 can lead to excessive collagen crosslinking, which can cause tissue remodeling and fibrosis (74). These findings suggest that ECM remodeling is not a linear process but involves multiple compensatory pathways aimed at preserving structural integrity and biomechanical properties of the TM.

To our surprise, among the top 10 upregulated proteins, we found elevated levels of ADM under siCTSK. A clinical study suggested higher aqueous humor ADM levels in patients with primary open-angle glaucoma than in those with neovascular glaucoma (75). In an earlier study (76), ADM was shown to lower IOP upon intravitreal injection in rabbits in a dose-dependent manner and interestingly, the C-terminal fragment of ADM (AM22-52) did not lower IOP. Additionally, ADM has been shown to upregulated under shear stress (77). ADM has been shown to elicit a dose-related increases in mean arterial pressure after cerebroventricular administration (78) suggesting a differential local and distal effects. Thus, suggesting ADM could be playing an important role in regulating IOP and a possible role of ADM either in pathogenesis or adaptative behavior of the TM cells/tissue. ADM has been implicated in ECM remodeling and oxidative stress adaptation (43, 44, 46). Given the extensive ECM changes observed in CTSK-depleted TM cells, ADM may be part of a broader regulatory mechanism aimed at maintaining TM tissue homeostasis. Notably, ADM is known to influence endothelial barrier function and fibrosis progression (79), raising the question of whether ADM signaling contributes to TM contractility and aqueous humor outflow regulation. Future studies are required to explore functional role of ADM in modulating ECM and cytoskeletal rearrangements in TM regulated by CTSK. Our data showed that MMP9 activation was significantly reduced in CTSK-depleted cells, as evidenced by zymography analysis, which aligns with previous reports indicating importance of CTSK in MMP9 activation (67). Since MMPs contribute to ECM degradation and turnover, reduced MMP9 activation may further exacerbate ECM dysregulation following CTSK loss.

Since ECM remodeling influences integrin-mediated signaling and mechanotransduction, we next examined its impact on intracellular calcium levels, a key regulator of TM contractility and cytoskeletal remodeling (80, 81). Calcium imaging assays revealed sustained increase in intracellular calcium levels following CTSK depletion. Interestingly, CTSK loss of function suggests a sustained increase in intracellular calcium levels. This increase was observed in both pharmacological CTSK inhibition and siRNA-mediated CTSK knockdown, suggesting that calcium homeostasis is affected independent of CTSK enzymatic activity. The mechanism underlying calcium dysregulation remains unclear but may involve ECM-integrin interactions and downstream signaling pathways. Interestingly, our proteomics analysis identified an upregulation of CXCL10, CXCL11, and ITPRIP, all of which have been implicated in intracellular calcium mobilization. CXCL10 and CXCL11 primarily signal through CXCR3 and ACKR3, receptors known to influence GPCR-mediated calcium signaling (40, 41), while ITPRIP interacts with inositol 1,4,5-trisphosphate receptor (IP3R) to regulate calcium release from the endoplasmic reticulum (42). These findings suggest that CTSK depletion may trigger a broader signaling cascade that disrupts intracellular calcium homeostasis. Given the increase in intracellular calcium and its role in cytoskeletal dynamics, we further investigated the impact of CTSK depletion on actin polymerization. Proteomics analysis revealed a significant upregulation of PRKD1, a key regulator of actin remodeling and focal adhesion assembly (22). Immunoblot analysis confirmed increased PRKD1 phosphorylation, supporting its potential activation.

PRKD1 is known to regulate actin polymerization through the LIMK1/SSH1/cofilin pathway (70, 82). Consistent with this, CTSK depletion resulted in increased phosphorylation of LIMK1 and SSH1, leading to cofilin inactivation through phosphorylation, enhanced actin polymerization and increased focal adhesion assembly in CTSK-deficient TM cells. Since PRKD1 activation has been linked to focal adhesion maturation (83), these findings suggest that CTSK depletion promotes cytoskeletal rearrangement and adhesion signaling through PRKD1-mediated mechanisms. Interestingly, while we observed an increase in actin polymerization and focal adhesion formation following CTSK depletion, several myosin motor proteins (MYH7, MYH6, MYH13) and RhoQ were significantly downregulated in our proteomics data. Myosin proteins are crucial for actin-myosin interactions that drive cellular contractility and mechanotransduction (52, 53, 84–86). The reduction in myosin under CTSK depletion could allow more free actin monomers to assemble into filaments without myosin to “use up” the actin, favoring increased actin polymerization. RhoQ is a small GTPase involved in actin cytoskeletal rearrangement (87). Interestingly, RhoQ knockdown has been reported to induce TGF-β-induced fibrosis in lung adenocarcinoma (88), suggesting a possible shift in cytoskeletal and ECM regulation following CTSK depletion. Since PRKD1 has been shown to regulate RhoA activation via Rhotekin phosphorylation and SSH1L-mediated signaling (74, 75), it remains unclear whether PRKD1 activation in our study influences broader Rho GTPase signaling. Further studies are required to determine whether PRKD1-mediated actin polymerization compensates for the loss of RhoQ and myosin-driven contractility.

Finally, the conundrum lies in increase in apoptotic markers, including CASP3, CASP7, TRADD, and PPM1F (47–49) without caspase 3/7 activation following CTSK knockdown may reflect a priming effect where TM cells are poised for apoptosis but do not fully undergo programmed cell death. Alternatively, these markers may have non-apoptotic roles in cellular stress responses (89–91).

The major limitation of this study is the choice of *in vitro* model to characterize the loss of CTSK function and future *in vivo* validation using CTSK knockout mice is necessary to confirm its physiological relevance. Additionally, investigating the functional consequences of changes mentioned above on TM biomechanics and aqueous humor outflow will be critical to understanding how CTSK loss impacts IOP regulation. The effect of CTSK loss on lysosomal function in TM also warrants further studies.

In conclusion, our findings suggest that CTSK plays an important role in maintaining ECM integrity and cytoskeletal regulation in TM cells. Its depletion leads to significant disruption in ECM homeostasis and actin polymerization (**Figure 5**). Given the role of TM dysfunction in glaucoma, targeting CTSK-related pathways may offer novel therapeutic strategies for regulating IOP and preventing disease progression.

## Author contributions

Conceptualization: PPP and AS. Methodology: Proteomics, cell cultures, immunofluorescence, immunoblotting, and functional assays: AS, KS, ED, RPP, MS and PPP. Formal analysis: AS, KS, ED, PPP. Investigation: AS and PPP. Data curation: AS and PPP. Writing—original draft preparation: AS, KS and PPP. Figure preparation: AS, KS and MS. Writing—review and editing: AS and PPP. Visualization: AS and PPP. Supervision: PPP. Project administration: PPP. Funding acquisition: PPP. All authors have read and agreed to the published version of the manuscript.

## Institutional Review Board statement

Ethical review and approval were waived for the use of TM cells from human cadaveric corneal rims for this study.

## Acknowledgments

We would like to acknowledge Dr. Emma Doud from the Center for Proteome Analysis, IUSM, for helping with LC-MS/MS, data analysis, data evaluation, verification, and methods write-up. This project was supported by the National Institutes of Health/National Eye Institute R01EY029320, R01EY035412, and R01EY036107 (PPP); Shaffer Grant from The Glaucoma Foundation (PPP); Award from the Ralph W. and Grace M. Showalter Research Trust and the Indiana University School of Medicine (PPP), Research Support Funds Grant (PPP), Cohen AMD Research Pilot Grant (PPP), RPB Departmental Pilot Grant (PPP), Glick Research Endowment Funds (PPP), and Challenge grant from Research to Prevent Blindness to IU.

---

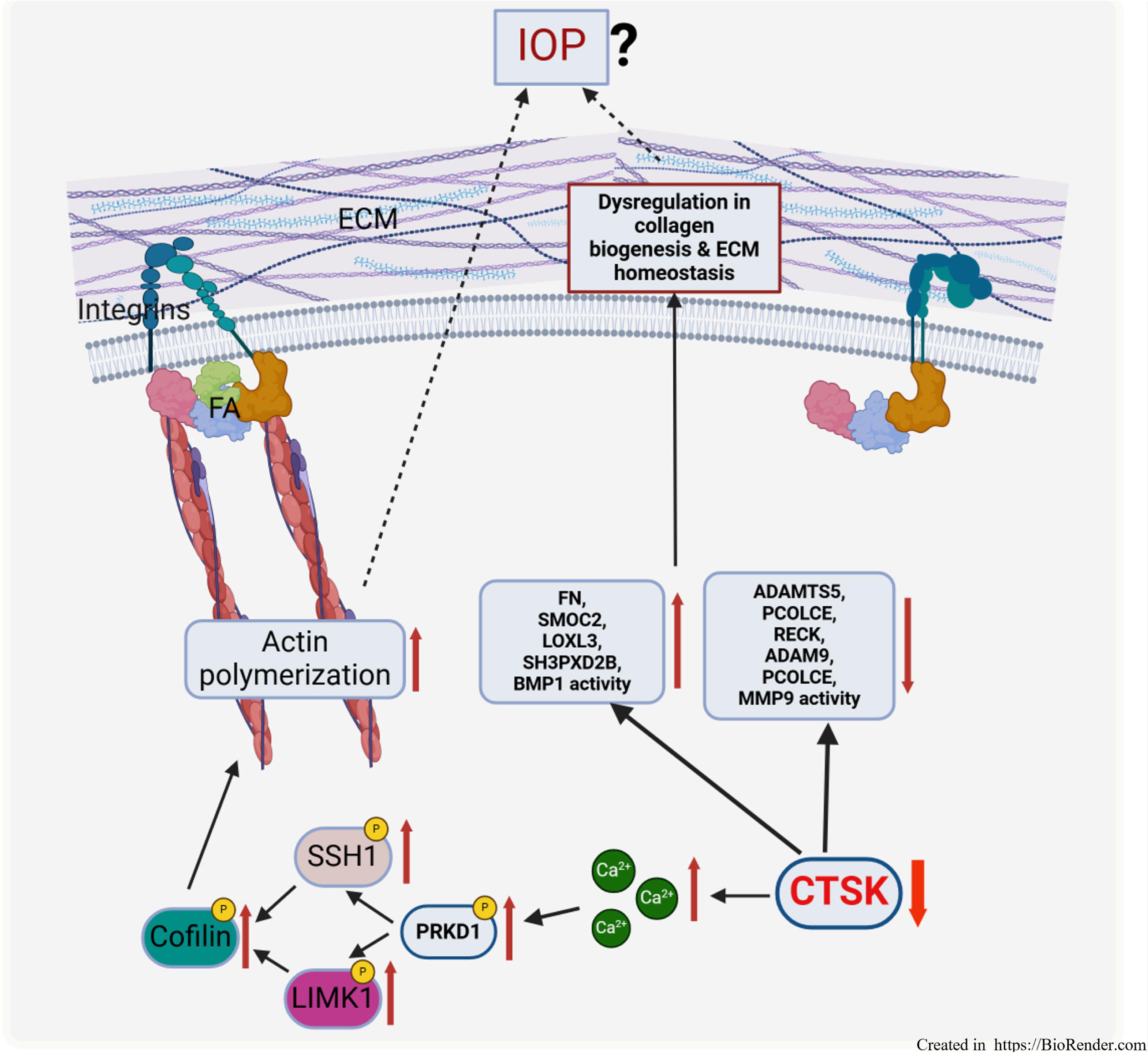

